# Root specific plasticity induced by paclobutrazol confers improved deficit irrigation tolerance and agronomic performance in maize

**DOI:** 10.1101/2020.05.12.087940

**Authors:** Mohammad Urfan, Haroon Rashid Hakla, Shubham Sharma, Manu Khajuria, Santosh B. Satbhai, Dhiraj Vyas, Sunil Bhougal, Narendra Singh Yadav, Sikander Pal

**Affiliations:** Plant Physiology Laboratory, Department of Botany, University of Jammu, Jammu-180006, India; Biodiversity and Applied Botany Division, CSIR-Indian Institute of Integrative Medicine, Canal Road, Jammu 180001, India; Department of Biological Sciences, Indian Institute of Science Education and Research (IISER), Mohali, SAS Nagar, Punjab 140406, India; Department of Statistics, University of Jammu, Jammu-180006, India; Department of Biological Sciences, University of Lethbridge, Lethbridge, AB 403587, Canada

**Keywords:** maize, root plasticity, root system architecture, deficit irrigation, cob yield, water budgeting, economic attributes, phenotypic plasticity

## Abstract

Drought stress in maize often results in poor growth and reduced yield. Antioxidants play vital role in management of abiotic stresses. Drought or water deficit are detrimental to young seedlings establishment and transition from vegetative to reproductive growth in maize. Paclobutrazol (PBZ) has been widely used to confer abiotic stress tolerance in plants, however, its impact on root developmental attributes in maize and their relevance in drought management are least understood. Comprehensive experiments over a three year period (2017-2019) under early deficit (EDI) and terminal deficit (TDI) irrigation with or without paclobutrazol (PBZ) were conducted on five maize varieties (DDKL, DKC-9144, PG-2475, PG-2320 and Bio-9621). Plant shoot and root growth kinetics, phenological changes and physiological perturbations including antioxidant profile coupled with molecular regulation of root traits, showed DKC-9144 as best variety in terms of plant fitness and reproductive performance under deficit irrigation. Root developmental rates were key contributors towards improved plant biomass and cob yield under deficit irrigation tolerance. Structural equation modelling (SEM) demonstrated specific contribution of different root types (crown, brace and seminal roots) in maize towards improving water use efficiency, cob yield and plant height. From SEM, seminal root surface area and root length are proposed desired traits to improve water use efficiency and cob yield in DKC-9144 under deficit irrigation. Bi-variate analyses of twenty key traits of plant fitness and agronomic importance showed a strong correlation (*r*) between root traits and improved growth performance and yield stability indices.

## 1.0 INTRODUCTION

Maize is mainly a rain fed crop in Indian sub-continent. Annual monsoon pattern showed disturbance consistently in recent years and has become a great concern not only for India, but also other countries in South Asia, where ~1.9 billion people live (Manivasagam and Nagarajan 2018; Parmar *et al*., 2019). Sustainable usage of irrigation resources is offered by deficit irrigation practices over last few years. Maize cultivation is synchronized with monsoon, and few days’ late monsoon arrival causes’ negative impact on growth of young maize plants (Thomas and Prasannakumar 2016; Manivasagam and Nagarajan 2018; Laitonjam *et al*., 2018; Parmar *et al*., 2019). Irregular rain patterns and climate change impact maize cultivation pan India (Thomas and Prasannakumar 2016; Laitonjam *et al*., 2018). Soil water shortage in months of June-September often causes drought conditions or soil water deficit, resulting in poor plant growth and establishment of young maize plants (early water deficit) and successful transition of vegetative stage to reproductive stage (Deng *et al*., 2018; Xu *et al*., 2018). Water deficit poses alarming threat to crop loss worldwide and may reduce average 50% yield (David 2014; Thatcher *et al*., 2016; Wang *et al*., 2018; Dowswell 2019; He *et al*., 2019; Menkir *et al*., 2020). Impact of water deficit is proven more critical during flowering time (terminal water deficit) and results in approximately 40-60% yield loss (David 2014; Thatcher *et al*., 2016; Wang *et al*., 2018; Dowswell 2019; He *et al*., 2019; Menkir *et al*., 2020).

Maize responses to water deficit or drought are multiple and inter-linked with reduced leaf surface area, leaf water potential, relative water content and transpiration rate (Begcy *et al*., 2019; Sah *et al*., 2020). Sunken stomata, lowered stomatal density and shorter stem and delayed in onset of tassel and ear formation occur under drought (Sah *et al*., 2020). Root adaptations include shallower root system or deep root system depending on soil water availability, reduction or increase in lateral roots on all the three different root types’ viz. crown roots (CRs), brace roots (BRs) and seminal roots (SRs) (Hund *et al*., 2009; Comas *et al*., 2013; Zhan *et al*., 2015;Sebastian *et al*., 2016; Borrás and Vitantonio 2018; Turc and Tardieu 2018; Klein *et al*., 2020). Physiological perturbations induced under drought include reduced photosynthesis, photochemical efficiency and net assimilation rate (NAR) (Pal *et al*., 2016; Mohan *et al*., 2019). Root traits and their prospective role in water deficit management in crop plants are least explored in maize. Root traits such as root biomass, root length, root density, root surface area and root depth contribute towards water stress avoidance (Hund *et al*., 2009; Comas *et al*., 2013; Zhan *et al*., 2015; Sebastian *et al*., 2016; Borrás and Vitantonio 2018; Turc and Tardieu 2018; Klein *et al*., 2020).

Climate change and its impact on soil water availability and rain patterns across world have posed a serious challenge to crop management and world food security (Seneviratne 2012; Lesk *et al*., 2016; Jägermeyr and Frieler 2018; Zhang *et al*., 2018; Sloat *et al*., 2020). In Indian sub-continent, rain water is the main source of maize irrigation (85% of total maize cultivation); maize cultivators are facing a severe challenge of drought posed by erratic monsoon arrival or rain period (Challinor *et al*., 2016; Murari *et al*., 2018; Zaveri *et al*., 2020). India ranked seventh (17,300,000 tonnes) in maize production, with tally topped by USA (333,010,910 tonnes) at world level in 2016-2017. Productivity of maize /ha is dismal low in India (2.54 million MT, 2016-17) compared to world average (5.82 MMT, 2016-17) and US (10.96 MMT, 2016-17), with dependence on rain fall (monsoon-south west and north east monsoon), salinity and other abiotic and biotic stresses as main reasons (Challinor *et al*., 2016; Murari *et al*., 2018; Zaveri *et al*., 2020). Extensive state-wise data analyses of maize production in India over past 10-15 years revealed interesting observations. Nearly 15% of area under maize cultivation is irrigated, leaving 85% rain fed (mainly monsoon) source of irrigation (Sharma and Mujumdar 2017; https://www.farmer.gov.in/M_cropstaticsmaize.aspx). It is obvious to have low maize productivity in Indian states having irregular or shorter spell of monsoon rains. Data analysis of 1999-2018 showed a positive correlation between numbers of drought affected districts (DADs) and maize productivity at National level (Supplementary Table 1 and 2, Supplementary Fig. S1). Deep analysis revealed irregular monsoon episodes at state levels as a causative factor for reductions in maize productivity (https://mausam.imd.gov.in/). For instance, reduction in monsoon rains (2013-14, 2014-15 and 2015-16) in Andhra Pradesh, with increase in number of DADs showed a linear relationship with reduction in maize productivity and yield, though area under cultivation increased over consecutive years (Supplementary Fig.S1 b). Similar observations were noted for Karnataka, Madhya Pradesh, Uttar Pradesh and Union territory of Jammu and Kashmir (Fig.S1 c-f). Improving maize productivity offers a new paradigm to increase farmers’ income compared to other crops cultivated in India (http://ficci.in/ficci-in-news-page.asp?nid=14261). Country would require 45 million metric tons of maize by year 2022 and to achieve this target, innovative ideas of improving maize productivity in rain fed areas demands more attention (http://ficci.in/ficci-in-news-page.asp?nid=14261).

Deficit irrigation both in terms of early deficit irrigation (EDI) and terminal deficit irrigation (TDI) offers an alternate method of conventional irrigation, where in a substantial amount of irrigation water could be saved by reducing evapotranspiration demand (EVTD) of plants with minimal effects on yield performance (Davies and Bennett 2015; Barker *et al*., 2019; Trout and Manning 2019).

Paclobutrazol (PBZ) is a plant growth regulator of the triazole family, known to induce inhibition of cell elongation and internode extension *via* negatively affecting gibberellins biosynthesis (Pal *et al*., 2016; Mohan *et al*., 2019). Plants treated with PBZ have been shown to produce elevated levels of abscisic acid, which is a proven anti stress hormone in plants. PBZ may also induce morphological modifications of leaves in terms of number and smaller stomatal pores and thicker leaves. In general, PBZ also enhances stem diameter, root density, root length and roots surface area to confer abiotic stress tolerances in plants (Pal *et al*., 2016; Mohan *et al*., 2019). Earlier studies showed positive impact of paclobutrazol (PBZ) on water use efficiency (WUE) of tomato and mulberry plants *via* increasing root surface area and bringing physiological changes in both upper and lower parts of tomato and mulberry under drought (Pal *et al*., 2016; Mohan *et al*., 2019; Sloat *et al*., 2020). Understanding root growth kinetics (root surface area, root length and root numbers) in maize and its impact on shoot growth kinetics and overall plant growth performance under drought or deficit irrigation is least explored. Structural equation modelling will explore the contribution of different root traits in making maize plants more efficient in WUE, water productivity (WP) with minimal effect on yield performance (cob yield) and farmers income. Current study investigated impact of early deficit irrigation (EDI, mimicked late arrival of monsoon) and terminal deficit irrigation (TDI, mimicked shorter monsoon spell) on overall plant fitness and yield performance in five maize varieties (DDKL, DKC-9144, PG-2475, PG-2320 and Bio-9621) over a three year period. Findings propose significant contributory role of root traits in making DKC-9144 as best adapted for deficit irrigation (EDI and TDI) with minimal impact on crop yield.

## 2.0 RESULTS

### 2.1 Paclobutrazol improves growth kinetics of maize plants under deficit irrigation

Early deficit irrigation (EDI) reduced shoot extension rate (SER) significantly in DDKL and DKC-9144 (at 35 days after sowing, DAS only), while increased SER in Bio-9621. EDI with paclobutrazol (PBZ) significantly improved SER in PG-2475 and PG-2320 at DAS 25 over EDI. Only PBZ reduced SER considerably in PG-2475 at 25 and 35 DAS over control (Fig. 1 a). In terminal deficit irrigation (TDI), significant reductions in shoot extension rate (SERt) in all varieties at 74 DAS and 84 DAS and for 94 DAS noted (except PG-2475) (Fig. 1b). PBZ with TDI notably reduced SERt at 64 DAS for DDKL. Improved SERt at 74 DAS noted for all varieties, 84 DAS (DDKL), compared to TDI alone recorded. Overall reduction in SERt was noted for PBZ in control conditions (except at 94 DAS), strikingly increased SERt noted for PG-2320 and Bio-9621 (Fig. 1 b). Visible differences in the growth of plants were noted under control and deficit irrigation conditions (Supplementary Fig. S2). Besides measuring shoot growth kinetics, shoot mophpometerics revealed impact of EDI and TDI on shoot height, stem thickness, leaf number and leaf surface area (only for TDI) with or without PBZ (Supplementary Results, Supplementary Fig. S3 panel A and panel B).

**Figure 1.**
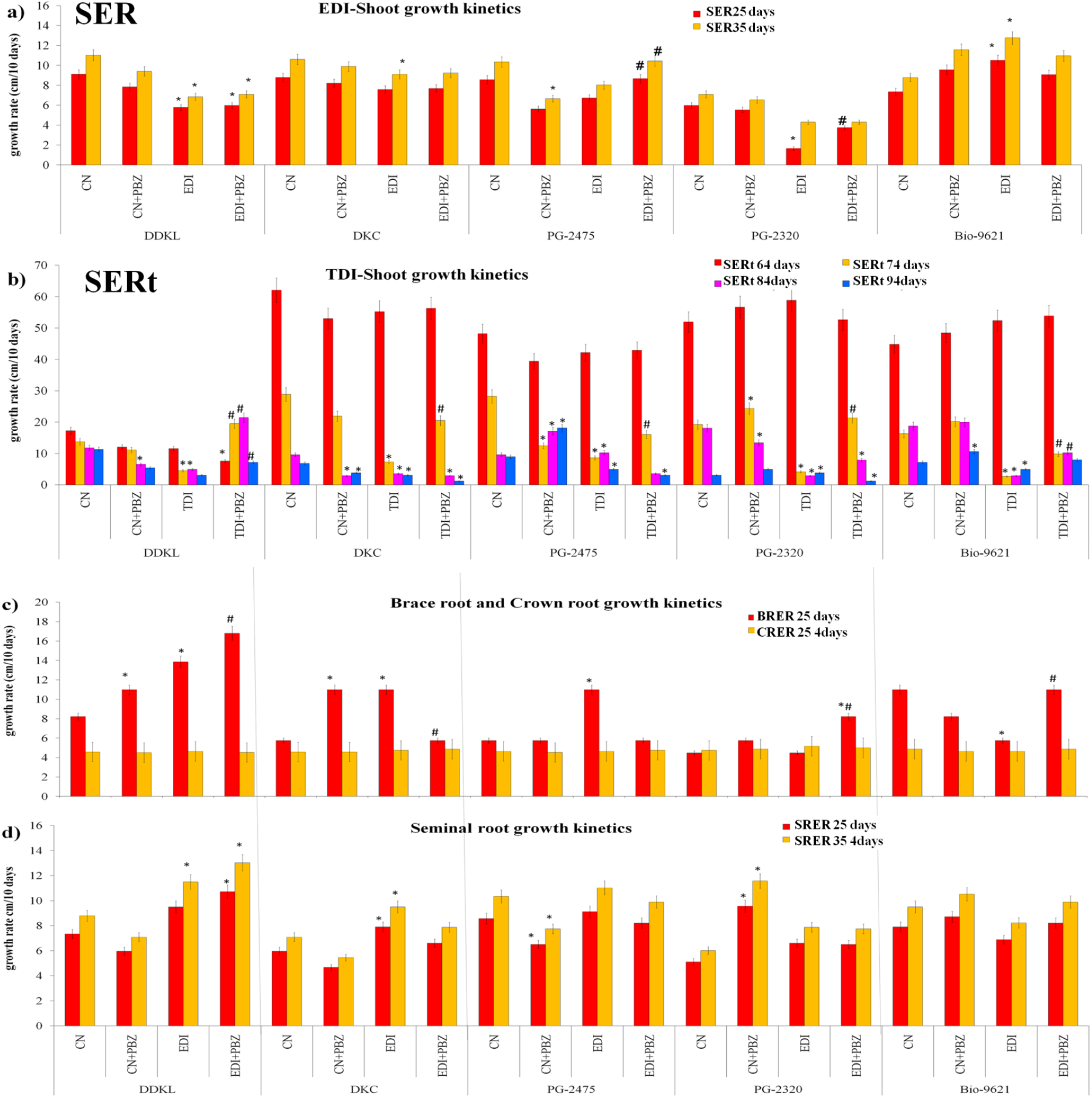
Paclobutrazol alter growth kinetics in maize to improve deficit irrigation tolerance. (a) Early deficit irrigation (EDI) imposed on young maize plants of DDKL, DKC-9144, PG-2475, PG-2320 and Bio-9621 at 15 days after sowing (DAS) until 35 DAS for a period of 20 days mimicked late arrival of rainwater spell of monsoon, altered shoot growth kinetics (a), showed EDI impact with or without paclobutrazol (PBZ) on shoot extension rate (SER, cm/10 days interval), such that SER 25 days calculated used 15 DAS shoot growth value as a base line, while SER 35 days calculated used 25 DAS shoot growth value as a base line. (b) Terminal deficit irrigation (TDI) imposed on maize plants of DDKL, DKC-9144, PG-2475, PG-2320 and Bio-9621 at 54 days after sowing (DAS) until 104 DAS for a period of 50 days mimicked shorter rainwater spell of monsoon, altered shoot growth kinetics measured at a gap of ten days each at 54-64, 64-74, 74-84, and 84-94, showed impact of TDI with or without paclobutrazol (PBZ) on shoot extension rate (SERt, cm/10 days interval) of maize plants, such that SERt 64 days calculated used 54 DAS shoot growth value as a base line, SERt 74 days calculated used 64 DAS shoot growth value as a base line and similar for 84 and 94 DAS SERt. (c-d) Early deficit irrigation (EDI) imposed on young maize plants of DDKL, DKC-9144, PG-2475, PG-2320 and Bio-9621 at 15 days after sowing (DAS) until 35 DAS for a period of 20 days mimicked late arrival of rainwater spell of monsoon, showed impact of EDI with or without paclobutrazol (PBZ) on root extension rate (RER, cm/10 days interval) of brace root (BR) and crown root (CR), seminal roots (SR) extension rates, such that RER 35 days of BR, CR was calculated with 25 DAS BR and CR growth value as a base line, (d) For SR, RER was calculated both at 25 DAS and 35 DAS, such that RER 25 days of SRs calculated used 15 DAS SRs growth value as base line, while SRER 35 days of SR calculated used 25 DAS SR growth value as a base line. Data presented are means ± standard errors (*n* = 15, biological replicates). Asterisk symbol (*) indicate significant differences from control in all combinations, while # indicate significant difference from EDI or TDI only (Tukey’s test, *P* ≤ 0.05). *Abbreviations:* control irrigation, CN, control+paclobutrazol CN+PBZ, early deficit irrigation, EDI, early deficit irrigation+paclobutrazol, EDI+PBZ, terminal deficit irrigation, TDI, terminal deficit irrigation+paclobutrazol, TDI+PBZ.

For root attributes, crown root extension rate (CRER) under EDI alone increased significantly in DDKL, DKC-9144 and PG-2475 over control. PBZ in EDI showed a significant increase in CRER for DDKL, PG-2475 and PG-2320 (Fig. 1c). PBZ in EDI or alone caused no significant changes in brace root extension rate (BRER) across all varieties (Fig. 1c). For seminal root extension rate (SRER) both at 25 and 35 DAS, EDI alone increased SRER in DDKL (35 DAS only) and DKC-9144 compared to control (Fig. 1d). PBZ with EDI at 25 and 35 DAS significantly increased SRER in DDKL over control (Fig. 1d). EDI and TDI alone showed visible reductions in root growth compared to control, while PBZ under EDI and TDI improved root growth over EDI and TDI alone (Fig. 2 a-b). In addition to root growth kinetics, laterals root number (LRNs) at 35 DAS (lateral roots on all root types viz. CR, BR and SR) were counted. Considerably improved LRNs under EDI compared to control were noted for DDKL, DKC-9144 and PG-2320 (Table 1). PBZ under EDI reduced LRNs significantly in DDKL, DKC-9144 and PG-2320. PBZ alone showed visible inhibitory effects on LRNs in all varieties expect DDKL (Table 1). PBZ was observed to modulate brace root, crown and seminal roots number and surface area under deficit irrigation (Supplementary Results, Supplementary Fig. S4). From shoot and root morphometeric observations under EDI and TDI alone with or without PBZ, DKC-9144 and DDKL emerged as best adapted varieties for deficit irrigation tolerance.

**Figure 2.**
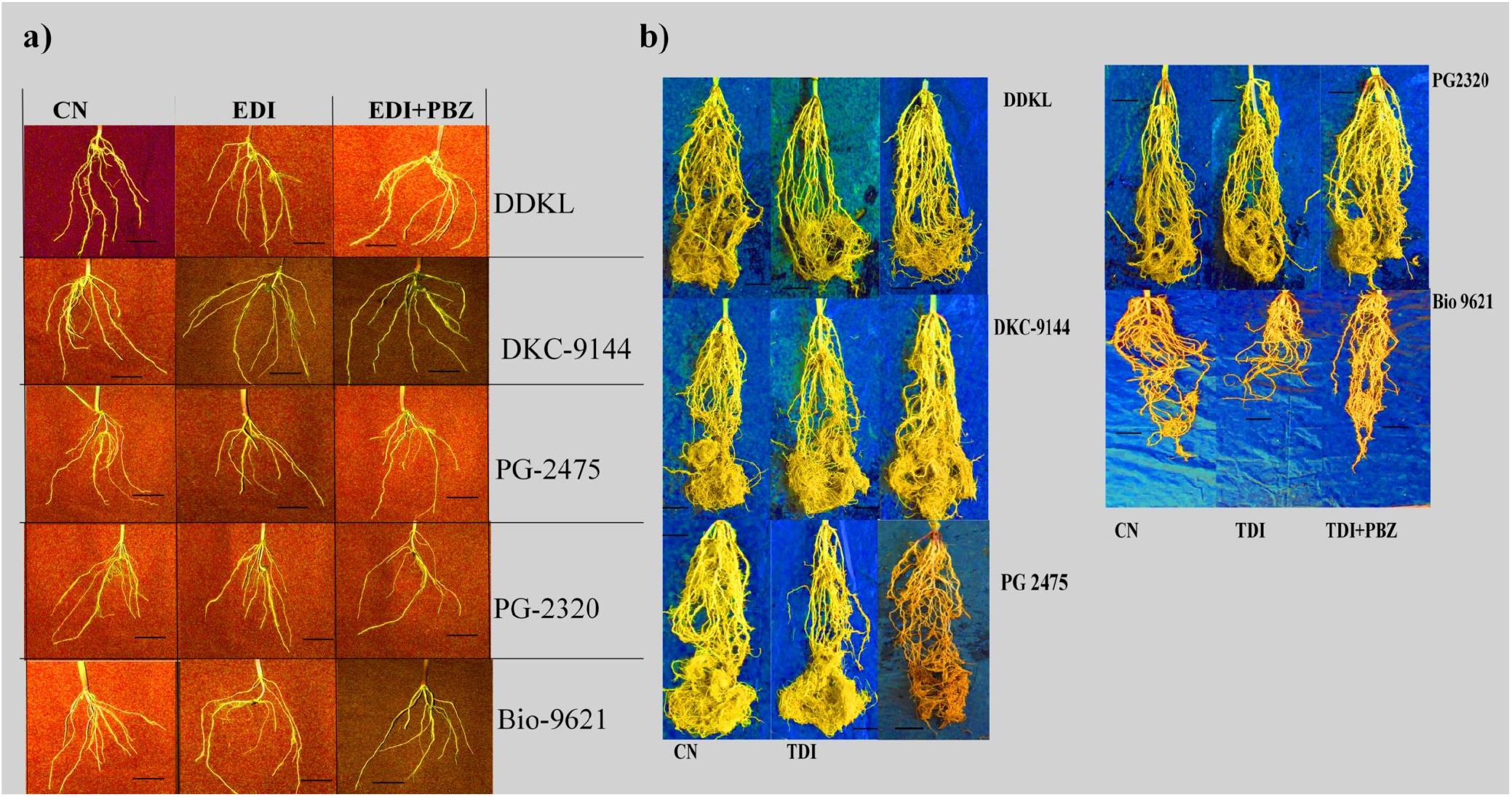
Paclobutrazol and deficit irrigation impacts root system of maize plants. (a) Harvested root system of young maize plants of DDKL, DKC-9144, PG-2475, PG-2320 and Bio-9621 at 35 days after sowing (DAS) subjected to early deficit irrigation (EDI, 60% of evapotranspiration demand, EVTD) with or without paclobutrazol (PBZ) at 15 DAS until 35 DAS for a period of 20 days mimicked late arrival of rainwater spell of monsoon, and (b) harvested root system of DDKL, DKC-9144, PG-2475, PG-2320 and Bio-9621 at 104 DAS subjected to terminal deficit irrigation (TDI, 60% of evapotranspiration demand, EVTD) with or without paclobutrazol (PBZ) at 54 DAS until 104 DAS for a period of 50 days mimicked shorter rainwater spell of monsoon. Scale bars represent 100 mm.

**Table 1.** Effect of early deficit irrigation (EDI) with or without paclobutrazol (PBZ) on lateral roots number (LRN) and terminal deficit irrigation (TDI) with or without PBZ on reproductive effort (RE%) for tassel formation (RE tassel %) and RE ear formation (RE ear %), leaf area index (LAI) and net assimilation rate (NAR g m^-2^ leaf area^-1^ °C day^-1^) rate of DDKL, DKC-9144, PG2475, PG2320 and Bio-9621 maize varieties subjected to control irrigation (100% evapotranspiration demand, EVTD), early deficit irrigation (60% EVTD) and terminal deficit irrigation (TDI, 60% EVTD) with or without paclobutrazol (60 ppm). Values indicate mean ± S.E, where number of biological replicates (*n* = 15). Different letters (a, b, c & d) in a column indicate significant differences from each other.

|  |  | LRN (EDI) | RE tassel % | RE ear% | LAI | NAR (g m<sup>-2</sup> leaf area<sup>-1</sup> °C d<sup>-1</sup>) |
| --- | --- | --- | --- | --- | --- | --- |
| **DDKL** | CN | 345±14a | 1 | 1 | 368±18 | 0.338±0.034 |
|  | CN+PBZ | 381±12a | 1.182 | 1.036 | 418±23b | 0.326±0.017 |
|  | TDI | 727±18b | 0.883 | 1.011 | 329±17 | 0.397±0.016a |
|  | TDI+PBZ | 352±16a | 1.441a | 1.005 | 340±13 | 0.394±0.013 |
| **DKC-9144** | CN | 723±29a | 1 | 1 | 400±26 | 0.334±0.019 |
|  | CN+PBZ | 455±8b | 0.991 | 0.978 | 461±8b | 0.298±0.004 |
|  | TDI | 1136±49c | 1.444a | 0.951 | 339±17a | 0.361±0.008a |
|  | TDI+PBZ | 456±34b | 1.336 | 0.947 | 366±11 | 0.386±0.014 |
| **PG-2475** | CN | 427±24a | 1 | 1 | 393±24 | 0.344±0.020 |
|  | CN+PBZ | 245±19b | 0.849 | 1.015 | 443±8b | 0.307±0.004 |
|  | TDI | 479±21a | 2.077a | 0.772a | 364±8 | 0.362±0.007 |
|  | TDI+PBZ | 435±27a | 0.546b | 1.421b | 380±16 | 0.351±0.013 |
| **PG-2320** | CN | 218±14a | 1 | 1 | 350±12 | 0.375±0.011 |
|  | CN+PBZ | 130±8b | 0.489a | 0.934 | 387±9 | 0.345±0.007 |
|  | TDI | 759±29c | 0.663 | 1.164 | 298±13a | 0.430±0.017a |
|  | TDI+PBZ | 1187±47d | 1.073 | 1.068 | 328±9 | 0.395±0.009 |
| **Bio-9621** | CN | 518±27a | 1 | 1 | 241±12 | 0.515±0.025 |
|  | CN+PBZ | 279±29b | 1.345 | 1.228a | 275±11 | 0.460±0.017 |
|  | TDI | 568±25a | 1.892a | 1.065 | 238±12 | 0.523±0.007 |
|  | TDI+PBZ | 457±21a | 0.378b | 1.136 | 249±15 | 0.535±0.024 |

### 2.2 Paclobutrazol improves reproductive traits of maize plants under deficit irrigation

TDI with or without PBZ influenced tassel (male inflorescence) developmental attributes, such that TDI alone increased diameter of tassel (diaT) in PG-2475 among all varieties (Fig. 3 a). Non significant increase in diaT in all varieties recorded for PBZ plus TDI over TDI. Length of tassel (LeT) under TDI increased in DDKL and decreased in DKC-9144 significantly over control (Fig. 3 b). PBZ with TDI increased LeT in DDKL and reduced in PG-2475, PG-2320 and Bio-9621. TDI reduced dry biomass of tassel (DBT) in DDKL, while increased it in PG-2475 and Bio-9621 significantly compared to control (Fig. 3 c). PBZ with TDI significantly increased DBT in DKC-9144 and PG-2320, while reduced DBT in PG-2475 and Bio-9621 over TDI. Reduced number of male flowers/plant and number of anthers/flower in DKC-9144, PG-2475 and Bio-9621 under TDI were noted over control (Fig. 3 d-e). PBZ plus TDI increased both number of male flowers/plant and number of anthers/flower significantly in DKC-9144 compared to TDI. Moreover, TDI considerably increased spikelet number/flower in all varieties except DKC-9144 over control (Fig. 3 f). PBZ with TDI reduced spikelet number/flower in all varieties except DKC-9144 and Bio-9621 over TDI (Fig. 3 f). Spikelet length under TDI reduced significantly in DDKL and DKC-9144, while increased it in PG-2475 significantly over control (Fig. 3 g). PBZ under TDI increased spikelet length significantly in PG-2475, PG-2320 and Bio-9621 over TDI alone. Silk length under TDI significantly reduced in DDKL, while increased in DKC-9144 over control (Fig. 3 h). PBZ under TDI improved silk length in all varieties except PG-2475 compared to TDI. Cob dry weight (CDW) in TDI alone reduced in all varieties significantly over control. PBZ under TDI increased CDW significantly in all varieties except DDKL compared to TDI (Fig. 3 i). Reproductive effort (RE%) for tassel formation increased in DKC-9144, PG-2475 and Bio-9621 under TDI over control. Further, RE% tassel increased in DDKL and PG-2320, while decreased in other varieties for PBZ plus TDI compared to TDI (Table 1). RE% ear under TDI reduced in PG-2475 over control, while PBZ under TDI, RE% ear maximally increased in PG-2475 compared to TDI (Table 1).

**Figure 3.**
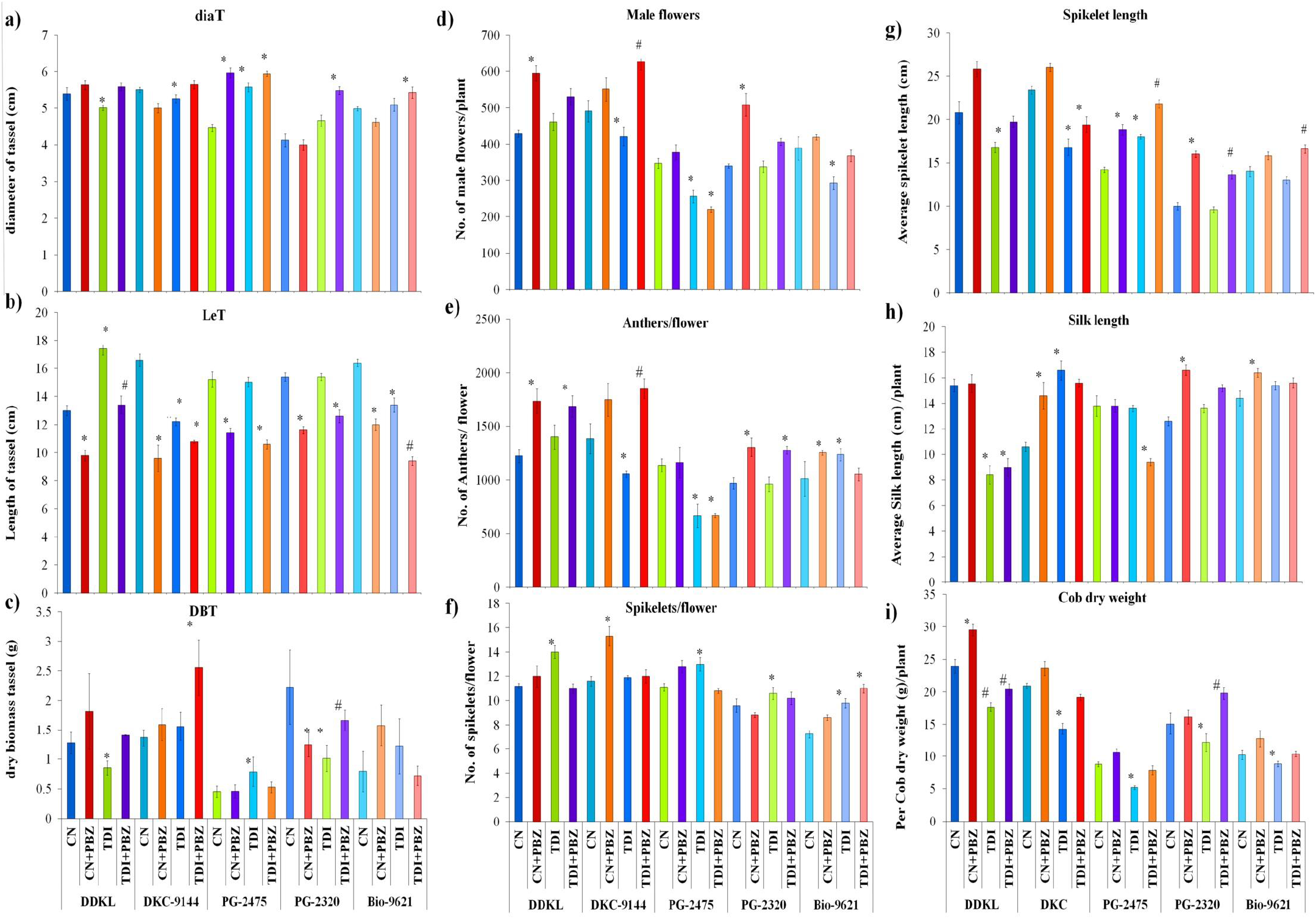
Paclobutrazol mitigates negative impacts of deficit irrigation on reproductive attributes of maize plants. Terminal deficit irrigation (TDI) with or without paclobutrazol (PBZ) imposed on maize plants of DDKL, DKC-9144, PG-2475, PG-2320 and Bio-9621 at 54 days after sowing (DAS) until 104 DAS for a period of 50 DAS mimicked shorter monsoon period, altered reproductive attributes. (a), tassel diameter (diaT, cm), (b) tassel length (LeT, cm), (c) dry biomass of tassel (DBT), (d) number of male flowers / plant, (e) average number of anthers /flower (anthers/flower), (f) number of spikelets/flower, (g) average spikelet length (cm), (h) silk length (cm) and agronomic trait viz. average cob dry weight (g)/plant at 104 DAS. Data presented are means ± standard errors (*n* = 10, biological replicates). Asterisk symbol (*) indicate significant differences from control in all combinations (Tukey’s test, *p* ≤ 0.05), while symbol # indicate significant difference from TDI irrigation in all combinations (Tukey’s test, *p* ≤ 0.05). *Abbreviations:* control irrigation, CN, control+paclobutrazol CN+PBZ, terminal deficit irrigation, TDI, terminal deficit irrigation+paclobutrazol, TDI+PBZ.

### 2.3 Plant biomass and water use efficiencies got improved by Paclobutrazol under deficit irrigation

For root dry biomass (RDB), TDI decreased brace roots dry biomass (BRDB), crown roots dry biomass (CRDB), seminal roots dry biomass (SRDB) and lateral roots dry biomass (LRDB) over control. PBZ plus TDI was seen to improve biomasses compared to TDI (Fig. 4 a-d). Under TDI, significant reduction in shoot dry biomass (ShDB) noted for DDKL, DKC-9144 and PG-2320 compared to control. PBZ under TDI improved ShDB in DKC-9144 and Bio-9621over TDI (Fig. 4 e). Similar pattern of total root dry biomass (TotRDB) was noted across all the varieties (Fig. 4f). For TDI alone, root/shoot ratio (RSR) significantly decreased in PG-2320 compared to control, while non significant changes occurred in other varieties. PBZ with TDI increased RSR in PG-2475 and Bio-9621 among all varieties over TDI. PBZ alone increased RSR in DKC-9144 and Bio-9621 over control (Fig. 4g). Evapotranspiration demand (EVTD) varied across all varieties at 54 DAS at the onset of TDI and PBZ application. EVTD declined under TDI in all maize varieties (Supplementary Fig. S5a). Enhanced EVTD in PBZ plus TDI in all varieties could be attributed to improved plant growth compared to TDI (Supplementary Fig. S5a). Water use efficiency (WUE) was measured for plant dry weight (PDW) basis (shoot + root) and CDW. TDI alone increased WUE-PDW significantly in all varieties except PG-2320 compared to control. PBZ under TDI improved WUE-PDW in DKC-9144, PG-2475 and PG-2320 compared to TDI significantly (Supplementary Fig. S5 b-c). PBZ under control increased WUE-PDW in all varieties in a non-significant manner. WUE-CDW under TDI significantly increased in DDKL, DKC-9144, and Bio-9621 over control (Supplementary Fig. S5 b-c). PBZ under TDI enhanced WUE-CDW in DDKL, PG-2475, and Bio-9621; while reduction noted in PG-2320 compared to control. PBZ alone increased WUE-CDW significantly in DDKL and Bio-9621 compared to control (Supplementary Fig. S5 b-c).

**Figure 4.**
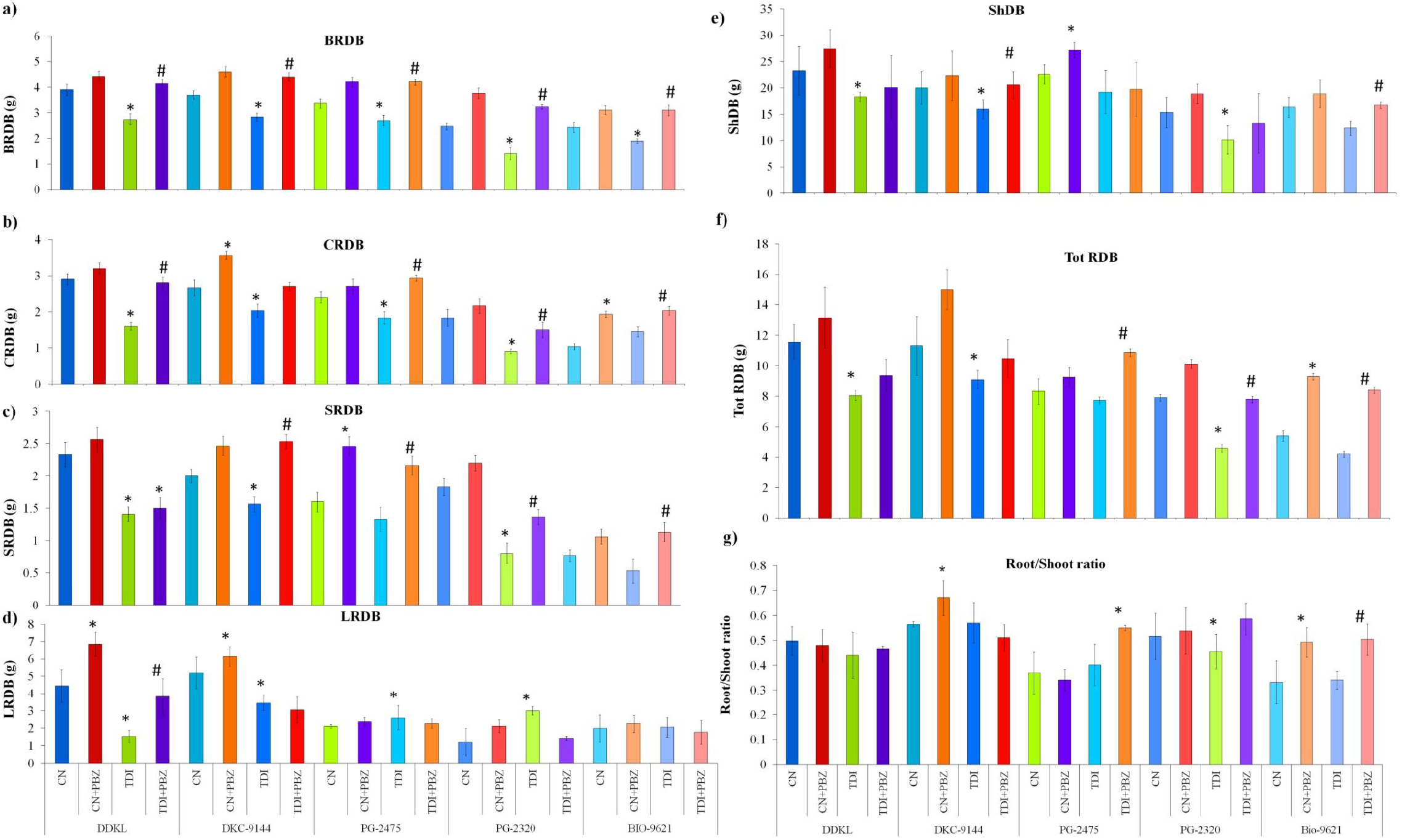
Paclobutrazol improves plant dry biomass under deficit irrigation. Terminal deficit irrigation (TDI) with or without paclobutrazol imposed on maize plants of DDKL, DKC-9144, PG-2475, PG-2320 and Bio-9621 at 54 days after sowing (DAS) until 104 DAS for a period of 50 DAS, showed impact on, (a) average brace root dry biomass (BRDB, g), (b) the average crown root dry biomass (CRDB, g), (c) average seminal root dry biomass (SRDB, g), (d) average lateral root dry biomass (LRDB, g), (e) the shoot dry biomass (ShDB), (f) total root dry biomass (TotRDB) and (g) the root/shoot ratio. Data presented are means ± standard errors (*n* = 15, biological replicates). Asterisk symbol (*) indicate significant differences from control in all combinations, while # indicate significant difference from TDI only (Tukey’s test, *p* ≤ 0.05). *Abbreviations:* control irrigation, CN, control+paclobutrazol CN+PBZ, terminal deficit irrigation, TDI, terminal deficit irrigation+paclobutrazol, TDI+PBZ

### 2.4 Paclobutrazol improves photosynthetic performance and modulates root architecture for deficit irrigation tolerance

Leaf area index (LAI) reduced in all varieties under TDI (Table 1). PBZ under TDI or alone non-significantly increased LAI in all maize varieties compared to TDI or control (Table 1). Net assimilation rate (NAR) under TDI was considerably increased in DDKL, DKC-9144 and PG-2320 over control. PBZ under TDI improved NAR in all varieties marginally. Non-significant reduction in NAR occurred for all varieties for PBZ alone (Table 1). Chlorophyll *a* and *b* contents under TDI declined in all varieties, whereas PBZ under TDI improved these pigments compared to TDI alone (Supplementary Fig. S6a-b).

Among five varieties, two varieties selected (on the basis of WUE-CDW and cob yield) viz. DKC-9144 (best performing) and PG-2475 (median performing) were examined for changes in photosynthetic parameters using LI-6400 XT infrared gas analyzer (Li-Cor, Lincoln, NE, USA) (Pal *et al*., 2016; Mohan *et al*., 2019). Net photosynthetic rate (P_N_) in EDI alone reduced in DKC-9144 and PG-2475 over control. PBZ under EDI at 35 DAS could improve P_N_ in DKC-9144 and PG-2475 over EDI (Fig. 5 a) in a non-significant manner. Intercellular CO_2_ (Ci) decreased significantly only at 25 DAS over control (Fig. 5 b). PBZ under EDI increased Ci in DKC-9144 and PG-2475 over EDI alone. Stomatal conductance (gs) increased in a non-significant manner under EDI alone and with PBZ at 25 DAS over control (Fig. 5 c). At 35 DAS EDI significantly reduced gs in DKC-9144 and PG-2475 over control; though small enhancements noted at 35 DAS for PBZ with EDI. Non-significant reductions in leaf transpiration rates (E) occurred in DKC-9144 and PG-2475 at 25 DAS under EDI alone. Contrastingly significant reductions in E occurred under EDI at 35 DAS in DKC-9144 and PG-2475 compared to control. PBZ under EDI at 35 DAS could improve E in a non significant manner in DKC-9144 and PG-2475 (Fig. 5 d). EDI alone increased total soluble sugars (sugar content) in DKC-9144 and PG-2475 over control (Fig. 5 e). Enhanced sugar contents were noted for PBZ under EDI in DKC-9144 andPG-2475 compared to EDI (Fig. 5 e). Improved photosynthetic parameters in DKC-9144 and PG-2475 in EDI plus PBZ could be linked with improvement in sugar content compared to EDI (Fig. 5 e).

**Figure 5.**
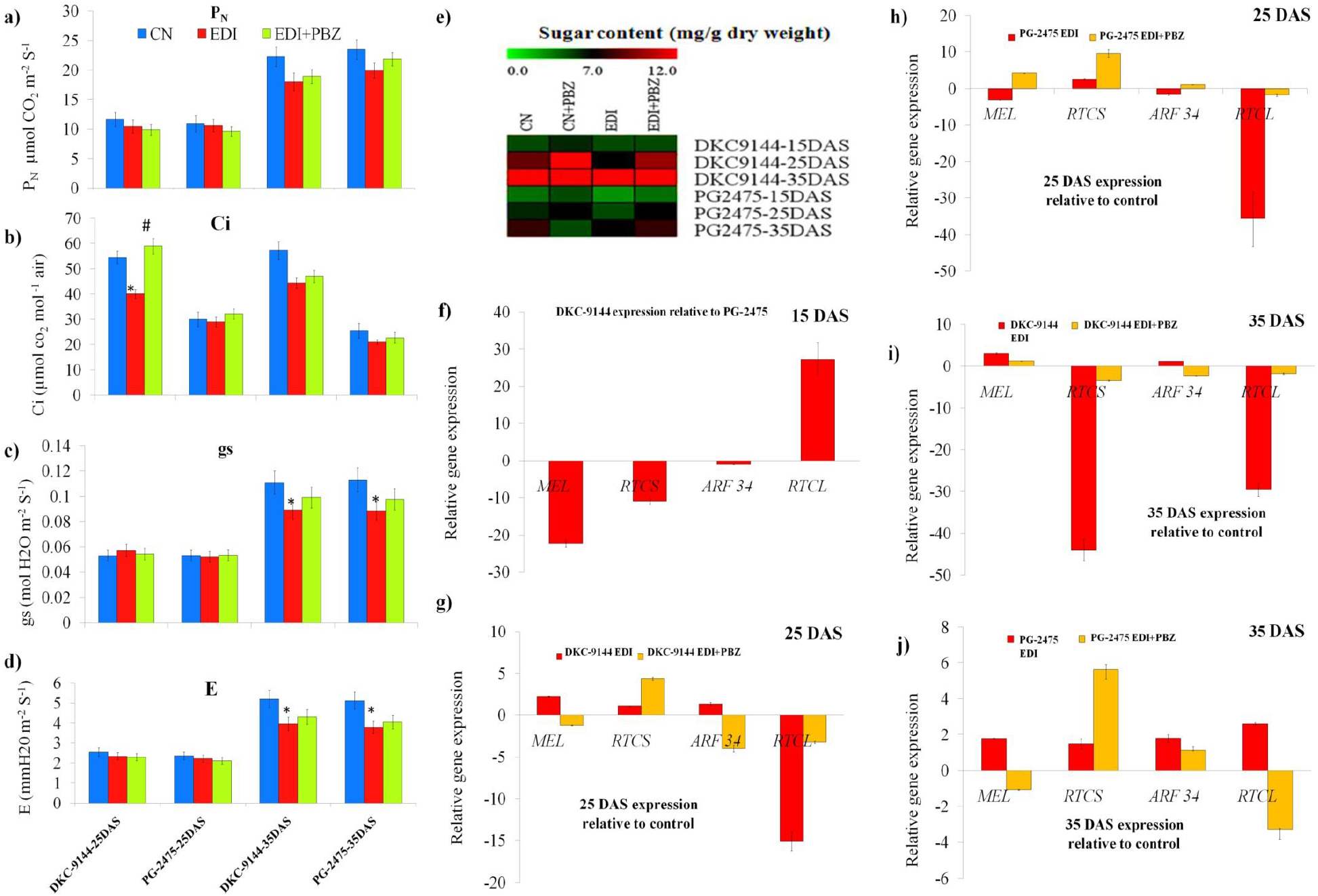
Paclobutrazol alleviates deficit irrigation impacts on photosynthetic efficiencies and molecular regulation of root growth in maize. Early deficit irrigation (EDI) with or without paclobutrazol (PBZ) imposed on young maize plants of DKC-9144 and PG-2475 at 15 days after sowing (DAS) until 35 DAS for a period of 20 DAS altered (a) net photosynthesis (P_N_, μmol CO2 m^-2^ S^-1^), (b) intercellular CO2 (Ci, μmol CO2 mol^-1^ air), (c) stomatal conductance (gs, mol H_2_O m^-1^ S^-1^) (d) transpiration rate (E, mm H_2_O m^-2^ S^-1^) (e) sugar content (mg/g dry weight), (f) relative expression of *ZmMel (metallothionein-like protein), ZmRTCS (gene regulating rootless crown and seminal roots), ZmRTCL* (*The RTCS-LIKE gene*), *ZmARF34* (*Zea mays Auxin response factor 34*) of DKC-9144 relative to PG-2475 (base expression) at 15 DAS before the commencement of EDI and PBZ application, (g-h) relative expression of *ZmMEL, ZmRTCS, ZmRTC* and, *ZmARF34* of DKC-9144 at 25 DAS after the commencement of EDI and PBZ in DKC-9144 relative to control (h) and of PG-2475 relative to control, (i-j) relative expression of *ZmMEL, ZmRTCS, ZmRTC* and, *ZmARF34* of DKC-9144 at 35 DAS after the commencement of EDI and paclobutrazol application in DKC-9144 (i) and of PG-2475 relative to control (j). Data presented are means ± standard errors (*n* = 15, biological replicates) for Figs a-d. Asterisk * and # indicate significant differences from control and early deficit irrigation respectively (Tukey’s test, *P* ≤ 0.05), Color code indicates differences across the row and column (for Fig. e). Data presented are means ± standard errors (*n* = 3, biological replicates, with each biological replicate had three technical replications) for Figs. f-j.

Relative expression of key genes regulating architecture of seminal and lateral root initiation was measured at different stages of EDI in DKC-9144 and PG-2475 (Fig. 5 f-j). At onset of experiment, (15 DAS), higher expression of *ZmRTCS, ZmARF34* and *ZmRTCL* was noted in DKC-9144 compared to PG-2475 (Fig. 5 f). At 25 DAS (10 days after deficit irrigation onset), EDI improved *ZmMel, ZmRTCS* and *ZmARF34* expressions, while significantly reduced expression of *ZmRTCL* in DKC-9144 compared to control (Fig. 5 h). However, in PG-2475 reduced expressions of *ZmMel, ZmARF34* and *ZmRTCL;* while elevated profiles of *ZmRTCS* were noted compared to control. PBZ under EDI reduced expressions of *ZmMel* and *ZmARF34* and *ZmRTCL*; while increased expression of *ZmRTCS* over EDI alone in DKC-9144. For PG-2475, PBZ with EDI improved expressions of *ZmMel, ZmRTCS, ZmARF34* and *ZmRTCL* compared to EDI alone (Fig. 5 h).

At 35 DAS (20 days after onset of deficit irrigation), EDI alone up-regulated expressions of *ZmMel, ZmARF34* and down-regulated *ZmRTCS* and *ZmRTCL* in DKC-9144; while in PG-2475 elevated expressions of *ZmMel, ZmRTCS, ZmARF34* and *ZmRTCL* compared to EDI alone noted (Fig. 5 i). PBZ under EDI at 35 DAS up-regulated *ZmRTCS* and *ZmRTCL* and down-regulated *ZmMel* and *ZmARF34* in DKC-9144 over EDI (Fig. 5 i). Under same conditions, PG-2475 showed elevated expressions of *ZmMel, ZmRTCS, ZmARF34* and *ZmRTCL* over control (Fig. 5 j). PBZ under EDI at 35 DAS showed down-regulation of *ZmMel* and *ZmARF34* and up-regulation of *ZmRTCS* and *ZmRTCL* compared to EDI alone (Fig. 5 j). Improved gene expressions of *ZmMel* and *ZmARF34* and down-regulated *ZmRTCS* and *ZmRTCL* in DKC-9144 could be linked with improved LRNs in DKC-9144 over PG-2475 (Table 1).

### 2.5 Paclobutrazol induced root plasticity enhanced deficit irrigation tolerance

Interesting behavioural pattern of different root types for plasticity indices was noted under deficit irrigation. Such that for EDI, maximum indices for BRs scope of plastic response (SPR) noted for DKC-9144, relative trait range (RTR) in Bio-9621, response coefficient (RC) and coefficient of variation (CoV) in DKC-9144, while for TDI maximum SPR and RTR for Bio-9621, RC for DDKL and CoV for PG-2475 (Table 2). For CRs under EDI and TDI, maximum SPR noted for DDKL and PG-2475, RTR for PG-2320 and PG-2475, RC for DDKL and DKC-9144 and CoV in DDKL and Bio-9621. For SRs, under EDI and TDI maximum SPR noted for DKC-9144 and PG-2320, RTR for PG-2320, RC for PG-2475 and DDKL, CoV for DKC-9144 and PG-2320 (Table 2). Visible changes in the root plasticity indices for BRs, CRs and SRs were noted under EDI and TDI when applied with PBZ compared to EDI and TDI alone (Table 2).

**Table 2.** Effect of early deficit irrigation (EDI, 60 % of evapotranspiration demand (EVTD)) and terminal deficit irrigation (TDI, 60% of EVTD) with or without paclobutrazol (PBZ) on the root plasticity indices (scope of plastic response, SPR), relative trait range (RTR), response coefficient (RC) and coefficient of variation (CoV) of brace roots, crown roots and seminal roots of DDKL, DKC-9144, PG2475, PG2320 and Bio-9621. Indices values outside brackets represent EDI with or without PBZ; while inside brackets represents TDI with or without PBZ. Number of biological replicates (*n* = 15).

|  |  | Scope of plastic response (SPR) | Relative trait range (RTR) | Response coefficient (RC) | Coefficient of variance (CoV) |
| --- | --- | --- | --- | --- | --- |
| Brace root | *CN/CN+PBZ-EDI (CN/CN+PBZ-TDI)* |  |  |  |  |
|  | DDKL | -0.5 (62.4) | -0.089 (0.060) | 0.918 (1.064) | 0.060 (0.030) |
|  | DKC-9144 | 0.1 (59.1) | 0.0163 (0.059) | 1.01 (1.063) | 0.011 (0.080) |
|  | PG-2475 | -0.3 (59.5) | -0.053 (0.062) | 0.949 (1.066) | 0.036 (0.088) |
|  | PG-2320 | 1.2 (50.9) | 0.1448 (0.064) | 1.169 (1.069) | 0.110 (0.044) |
|  | Bio-9621 | -0.7 (56) | -0.101 (0.075) | 0.907 (1.081) | 0.068 (0.028) |
|  | *CN/EDI (CN/TDI)* |  |  |  |  |
|  | DDKL | 0.2 (78.9) | 0.032 (0.081) | 0.967 (0.919) | 0.023 (0.081) |
|  | DKC-9144 | 1.2 (89.9) | -0.2 (0.096) | 1.2 (0.904) | 0.128 (0.030) |
|  | PG-2475 | 0 (133.8) | 0 (0.148) | 1 (0.851) | 0 (0.110) |
|  | PG-2320 | 0.3 (100.2) | 0.042 (0.135) | 0.957 (0.864) | 0.030 (0.088) |
|  | Bio-9621 | 0.9 (169.3) | 0.118 (0.246) | 0.881 (0.753) | 0.089 (0.083) |
|  | *EDI/EDI+PBZ (TDI/TDI+PBZ)* |  |  |  |  |
|  | DDKL | -0.8 (33.2) | -0.156 (0.035) | 0.864 (1.037) | 0.102 (0.055) |
|  | DKC-9144 | -1.3 (112.8) | -0.220 (0.117) | 0.819 (1.133) | 0.140 (0.094) |
|  | PG-2475 | 0.9 (63.7) | 0.132 (0.076) | 1.152 (1.083) | 0.100 (0.036) |
|  | PG-2320 | 1 (83.2) | 0.128 (0.115) | 1.147 (1.130) | 0.096 (0.072) |
|  | Bio-9621 | 1.1 (56.6) | 0.141 (0.098) | 1.164 (1.109) | 0.107 (0.014) |
| Crown root | *CN/CN+PBZ-EDI (CN/CN+PBZ-TDI)* |  |  |  |  |
|  | DDKL | 0.7 (15.6) | 0.104 (0.041) | 1.116 (1.043) | 0.077 (0.044) |
|  | DKC-9144 | 0.7 (51.7) | 0.095 (0.108) | 1.106 (1.121) | 0.071 (0.043) |
|  | PG-2475 | 0 (57.5) | 0 (0.117) | 1 (1.133) | 0 (0.045) |
|  | PG-2320 | 0.6 (19) | 0.082 (0.060) | 1.089 (1.064) | 0.060 (0.047) |
|  | Bio-9621 | 0.6 (11.5) | 0.088 (0.039) | 1.096 (1.041) | 0.065 (0.055) |
|  | *CN/EDI (CN/TDI)* |  |  |  |  |
|  | DDKL | 1.2 (39.1) | -0.2 (0.108) | 1.2 (0.891) | 0.128 (0.059) |
|  | DKC-9144 | 0.1 (18) | -0.015 (0.042) | 1.015 (0.957) | 0.010 (0.071) |
|  | PG-2475 | 0.3 (62.5) | -0.049 (0.144) | 1.049 (0.855) | 0.033 (0.113) |
|  | PG-2320 | 0.2 (35) | 0.029 (0.117) | 0.970 (0.882) | 0.021 (0.102) |
|  | Bio-9621 | 1 (31) | -0.161 (0.110) | 1.161 (0.889) | 0.105 (0.199) |
|  | *EDI/EDI+PBZ (TDI/TDI+PBZ)* |  |  |  |  |
|  | DDKL | -0.3 (26.1) | 0.043 (0.075) | 0.958 (1.081) | 0.030 (0.026) |
|  | DKC-9144 | 0.3 (58.1) | 0.042 (0.124) | 1.044 (1.142) | 0.032 (0.088) |
|  | PG-2475 | 0.1 (19.7) | 0.015 (0.050) | 1.015 (1.053) | 0.010 (0.056) |
|  | PG-2320 | 0.6 (28.5) | 0.084 (0.098) | 1.092 (1.108) | 0.062 (0.086) |
|  | Bio-9621 | -0.6 (-4.9) | -0.090 (-0.020) | 0.916 (0.980) | 0.061 (0.073) |
| Seminal root | *CN/CN+PBZ-EDI (CN/CN+PBZ-TDI)* |  |  |  |  |
|  | DDKL | 0.1 (78.1) | 0.015 (0.218) | 1.015 (1.28) | 0.010 (0.174) |
|  | DKC-9144 | 1 (71.8) | 0.142 (0.207) | 1.166 (1.261) | 0.108 (0.163) |
|  | PG-2475 | -0.2 (25.5) | -0.034 (0.070) | 0.966 (1.076) | 0.023 (0.052) |
|  | PG-2320 | 0.1 (25.3) | 0.016 (0.067) | 1.016 (1.072) | 0.011 (0.049) |
|  | Bio-9621 | -0.3 (18.9) | -0.056 (0.068) | 0.946 (1.074) | 0.038 (0.050) |
|  | *CN/EDI (CN/TDI)* |  |  |  |  |
|  | DDKL | 0.2 (-18.2) | -0.031 (-0.065) | 1.031 (1.065) | 0.021 (0.045) |
|  | DKC-9144 | 1.4 (8.7) | -0.233 (0.031) | 1.233 (0.968) | 0.149 (0.022) |
|  | PG-2475 | 1.4 (-0.7) | -0.233 (-0.002) | 1.263 (1.002) | 0.147 (0.001) |
|  | PG-2320 | 0.2 (47.2) | 0.033 (0.134) | 0.966 (0.865) | 0.024 (0.101) |
|  | Bio-9621 | 0.5 (15.2) | -0.089 (0.059) | 1.089 (0.940) | 0.060 (0.043) |
|  | *EDI/EDI+PBZ (TDI/TDI+PBZ)* |  |  |  |  |
|  | DDKL | -1 (19) | -0.178 (0.060) | 0.848 (1.063) | 0.115 (0.044) |
|  | DKC-9144 | -1 (20.4) | -0.156 (0.071) | 0.864 (1.076) | 0.102 (0.052) |
|  | PG-2475 | -1.6 (11.9) | -0.275 (0.034) | 0.783 (1.035) | 0.171 (0.024) |
|  | PG-2320 | 2.4 (30.5) | 0.296 (0.091) | 1.421 (1.100) | 0.245 (0.067) |
|  | Bio-9621 | -0.4 (5.7) | -0.070 (0.023) | 0.934 (1.023) | 0.047 (0.016) |

From drought susceptibility indices, most TDI susceptible variety noted was PG-2475 with stress susceptibility index (SSI); while PBZ under TDI reduced SSI maximally in DKC-9144 compared to TDI (Table 3). Maximum drought tolerance (TOL) under TDI recorded for DDKL, though PBZ improved TOL in all other varieties too. Mean productivity (MP) under TDI was higher in DDKL followed by DKC-9144; while PBZ improved MP significantly in DDKL and DKC-9144 over. Geometric mean productivity (GMP) was higher for TDI in DDKL and DKC-9144; while PBZ under TDI improved GMP in DDKL and DKC-9144 over TDI significantly (Table 3). Maximum stress tolerance index (STI) was noted for DDKL and DKC-9144 in TDI alone; PBZ under TDI improved STI in DDKL and DKC-9144 significantly. Maximum yield stability index (YSI) under TDI noted for DDKL and DKC-9144, furthermore, PBZ improved YSI in all varieties compared to TDI in a non-significant manner. Significantly maximum yield index (YI) observed for DDKL and DKC-9144 under TDI; while PBZ under TDI improved YI in DDKL and DKC-9144.For TDI alone, maximum harmonic mean (HM) was noted for DDKL and DKC-9144, while PBZ improved HM significantly in DDKL and DKC-9144 compared to TDI (Table 3).

**Table 3.**
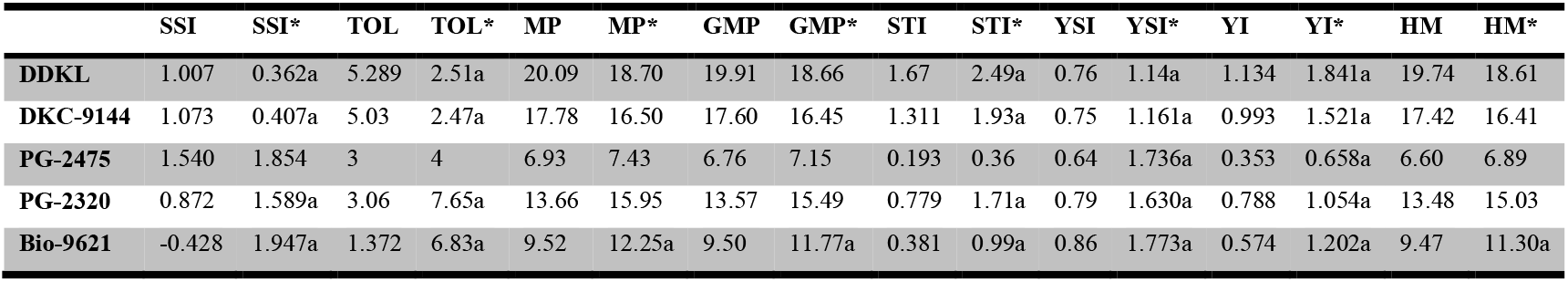
Performance of drought related indices, stress susceptibility index (SSI), tolerance (TOL), mean productivity (MP), geometric mean productivity (GMP), stress tolerance index (STI), yield stability index (YSI), yield index (YI) and harmonic mean (HM) of DDKL, DKC-9144, PG-2475, PG-2320 and Bio-9621 varieties subjected to control irrigation (100% evapotranspiration demand, EVTD), terminal deficit irrigation (TDI, 60% EVTD) with or without paclobutrazol (PBZ, 60 ppm). Values indicate mean ± S.E, where number of biological replicates (*n* = 15). Values without asterisk indicate ratio of cob yield under TDI over control, while asterisk symbol (*) indicate ratio of paclobutrazol plus TDI over TDI. Letter *a* row wise indicate significant difference between values of TDI vs PBZ plus TDI.

### 2.6 Structural equation modelling revealed root traits associated with water use efficiencies and growth performances

In control conditions in EDI experiment, structural equation modelling (SEM) showed differential impact of different root types (CR, BR and SR) on shoot height (SH), stem thickness (ST) and water use efficiencies (WUE) among DKC-9144 and PG-2475 (Fig. 6 a-b). For EDI alone in DKC-9144 over PG-2475, the effect of CR and BR was positive on SH than WUE and negative on ST (Fig. 6 c-d). SR impact was positive on ST and WUE but negative on SH for DKC-9144 over PG-2475 (Fig. 6 c-d). Under EDI plus PBZ among all root types, effect of SR on WUE, SH, and ST was more evident in DKC-9144 and PG-2475 (Fig. 6 e-f).

**Figure 6.**
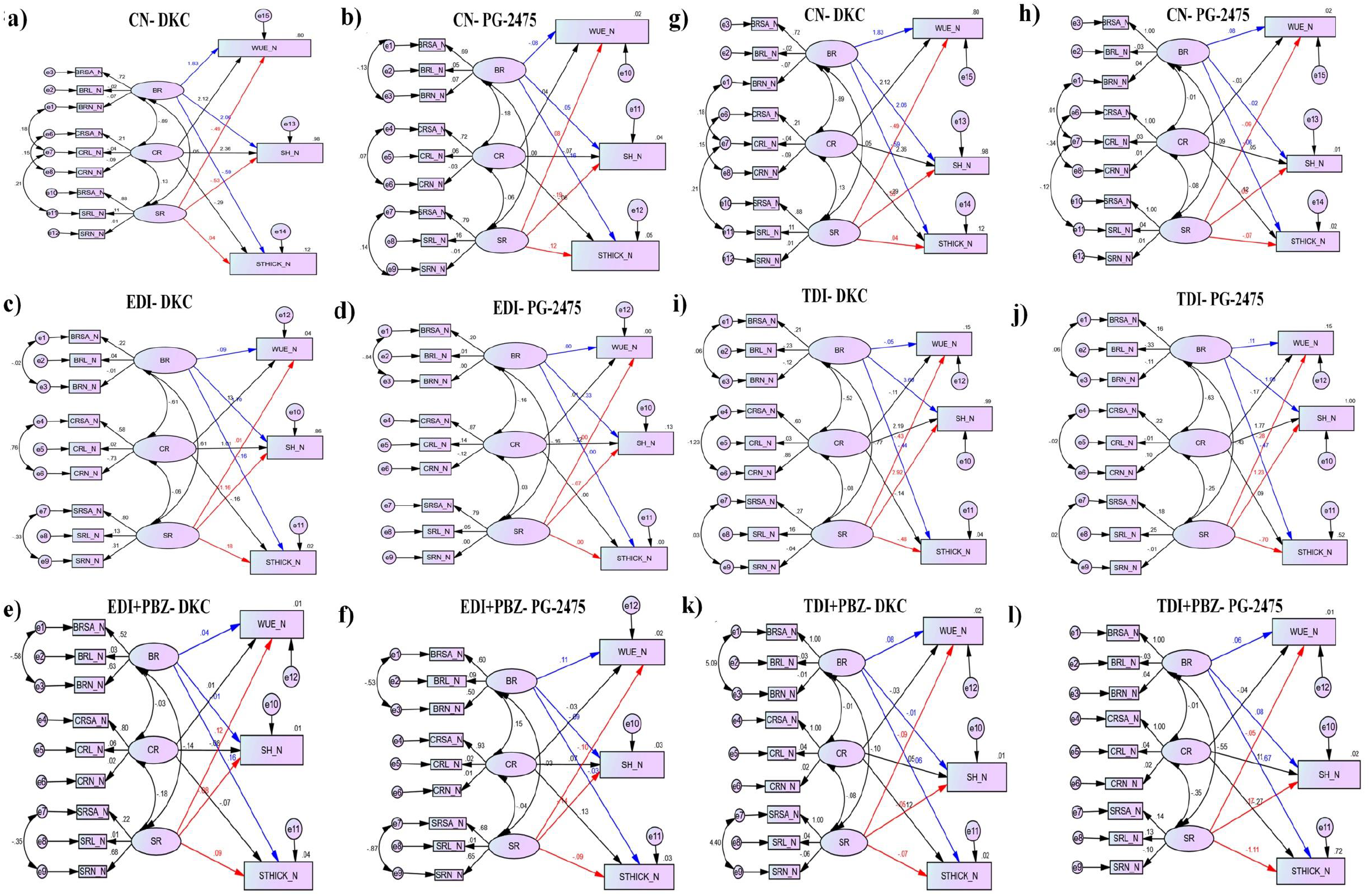
Structural equation modelling of maize root traits and their contribution in water use efficiency and plant fitness. Structural equation modelling (SEM) performed on the root traits of maize plants subjected to early deficit irrigation (EDI) and terminal deficit irrigation (TDI) showed contribution of selected root factors towards improving water use efficiency (WUE-cob dry weight), shoot height (SH) and stem thickness (ST). (a-b) Under control irrigation regime (CN), SEM model showed contribution of different root types such as crown roots (CRs), brace roots (BRs) and seminal roots (SRs) and their different traits such root number (RN), root length (RL) and root surface area (RSA) towards WUE, SH and ST in DKC-9144 (a) and PG-2475 (b). (c-d) Under EDI (60% of evapotranspiration demand), SEM model showed contribution of different root types such as CRs, BRs and SRs and their different traits such RN, RL and RSA towards WUE, SH and ST in DKC-9144 (c) and PG-2475 (d). (e-f) Under application of paclobutrazol (PBZ, 60 ppm) in EDI (EDI, 60% of evapotranspiration demand), SEM model showed contribution of different root types such as CRs, BRs and SRs and their different traits such RN, RL and RSA towards WUE, SH and ST in DKC-9144 (e) and PG-2475 (f). (g-h) Under control irrigation regime (CN) in TDI, SEM model showed contribution of different root types such as crown roots (CRs), brace roots (BRs) and seminal roots (SRs) and their different traits such root number (RN), root length (RL) and root surface area (RSA) towards WUE, SH and ST in DKC-9144 (a) and PG-2475 (b). (i-j) Under TDI (60% of evapotranspiration demand), SEM model showed contribution of different root types such as CRs, BRs and SRs and their different traits such RN, RL and RSA towards WUE, SH and ST in DKC-9144 (c) and PG-2475 (d). (k-l) Under application of paclobutrazol (PBZ, 60 ppm) in TDI (TDI, 60% of evapotranspiration demand), SEM model showed contribution of different root types such as CRs, BRs and SRs and their different traits such RN, RL and RSA towards WUE, SH and ST in DKC-9144 (e) and PG-2475 (f). **Blue, black and red** colored lines represents BRs, CRs and SRs, contributions of these root traits i.e. RN, RL and RSA for WUE, SH and ST are given in scores placed over respective colored lines merging into WUE, SH and ST.

For TDI under control conditions, among all root types, CR showed positive impact on SH and WUE while negative for ST in both DKC-9144 and PG-2475 (Fig. 6 g-h). In TDI alone, in DKC-9144 over PG-2475, effect of BR and CR was negative on WUE and ST, while positive on SH (Fig. 6 i-j). The SR showed negative effects on WUE and ST, and positive on SH in DKC-9144, while the effect of SR on WUE and ST was negative and on SH was positive (Fig. 6 i-j). For TDI plus PBZ conditions, in

DKC-9144, the effect of BR on WUE and ST was positive, while on SH was negative. For PG-2475, effect of BR on WUE, SH and ST was positive (Fig. 6 k-l). Effect of CR on WUE was negative, while positive on SH and ST in DKC-9144. For PG-2475, effect of BR on WUE and ST was negative and for SH was positive. For DKC-9144, effect of SR on WUE, SH and ST was negative, while the effect of SR on WUE was positive and negative for SH and ST in PG-2475 (Fig. 6 k-l).

## 3.0 DISCUSSION

Morphological adaptations such as reduced shoot and root growth, leaf area, elongation rate and final leaf length have been reported in maize under drought (Avramova *et al*., 2016). Present study showed altered shoot and root growth kinetics and shoot and root morphometerics in all maize varieties under deficit irrigation (EDI and TDI) and paclobutrazol (PBZ) (Figs. 1 and 2, Supplementary Figs. S2, S3). Inter-varietal impact of EDI and TDI was least for DKC-9144, in particular with PBZ (Fig. 1) Flowering and early seed developments are sensitive to drought (Westgate 1994). Water deficient conditions have shown to lower water potential in both vegetative and reproductive sinks resulting in poor seed formation and grain filling (Li *et al*., 2018). A week-long water deficit could impact floral development, ear length and grain number in maize variety DH4866 (Zhuang *et al*., 2007). Significant reduction in silk growth rate and silk emergence in maize under water deficit conditions has been reported (Oury *et al*., 2016). From current literature, it is observed that yield reduction in drought or water deficit could be a selective strategy of young reproductive organs to abort, in order to strengthen older fertilized ovaries to mature and lead to ear formation (Turc and Tardieu 2018). Present study has shown significant negative impact of TDI on reproductive attributes of tassel and silk including tassel length, tassel diameter, silk length and ear formation and reproductive effort for tassel and ear formation, plant biomass and its mitigation upon PBZ in all varieties with best performance being noted for DKC-9144 (Figs. 3-4, Table 1).

Deep rooting is a strategy for drought avoidance in plants, accompanied by increased number of lateral roots in upper part of the soil is positively correlated with transpiration, shoot dry weight and stomatal conductance (Hund *et al*., 2009). Furthermore, reduced LRs number and branching is associated with improved deep rooting, leaf relative water content, stomatal conductance and shoot biomass. Recently, a positive correlation between SRs length and drought tolerance index has been noted in maize seedlings (Guo *et al*., 2020). Another study conducted on 22 maize cultivars subjected to water stress at flowering and grain filling stages showed a strong correlation between root traits and drought management (Al-Naggar *et al*., 2019). Current study showed enhanced LRN under TDI conditions in all maize varieties; while PBZ application under TDI reduced LRN considerably among all varieties except PG-2320 (Table 1). From observations made, PBZ induced reduction in LRNs may be associated with improved management of deficit irrigation tolerance in maize varieties.

Root plasticity in a plant is guided by anatomical features of roots, which is a genetically controlled attribute in maize (Schneider *et al*., 2020). Effective drought and deficit irrigation management by plants largely depends on root plasticity i.e. adaptive responses of roots in response to low water (Cai *et al*., 2018). Highly plastic root system has been shown to counter drought more effectively compared to a plant having less plastic root system (Pires *et al*., 2020). Thus usage of plasticity indices acknowledges better drought management in plants (Cai *et al*., 2018; Pires *et al*., 2020). Current study also showed a positive correlation between SPR, RTR, RC and CoV of BR, CR and SR (Table 2) for TDI with or without PBZ (Table 2). Inter-varietal drought indices showed better performances in DKC-9144 and DDKL under TDI or EDI, with effect being more evident when TDI or EDI were accompanied by PBZ (Table 2). Performance of drought related indices of a crop plant infers its quality management of drought induced stress and its impact on economic attributes such as ear formation, height and yield (Barutcular *et al*., 2016). Inferior values of drought indices viz. stress susceptibility index (SSI), tolerance (TOL), mean productivity (MP), geometric mean productivity (GMP), stress tolerance index (STI), yield stability index (YSI), yield index (YI) and harmonic mean (HM) have often been linked to poor performance of a crop under drought, while superior values deduce better performance (Barutcular *et al*., 2016). Present study have shown significant impact of PBZ on TDI induced drought indices, such that PBZ application was noted to improve overall performance of maize plant in terms of plant height and yield attributes (Table 3). Root traits such as surface area and root length have been linked with drought management in plants (Cai *et al*., 2017). Current study also offered a positive correlation between root traits and deficit irrigation tolerance, such that reduction in brace root surface area (BRSA) and seminal roots surface area (SRSA) and enhancement in crown roots surface area (CRSA) was noted in PBZ plus TDI compared to TDI alone (Supplementary Table 4). Improved brace root length (BRL), crown root length (CRL) and seminal root length (SRL) in PBZ plus TDI over TDI alone could be accounted for better deficit irrigation tolerance in all maize varieties with best performance being noted for DKC-9144 and DDKL (Supplementary Fig. S4 and Supplementary Table 4). Root architecture plays a pivotal role in water management in plants, such that specific initiation of CRs and SRs in maize is regulated by *ZmRTCS* and *ZmRTCL* and their binding patterns *ZmARF34* and *ZmMel* (Xu *et al*., 2015). Altered root architecture in DKC-9144 and PG-2475 under EDI and EDI plus PBZ could be linked with differential expression of genes (*ZmMel, ZmRTCS, ZmRTCL* and *ZmARF34*) regulating architecture of CRs and SRs (Fig. 5 f-j). Thus improved root system of DKC-9144 over PG-2475 could be accounted for better deficit irrigation tolerance than PG-2475.

Drought induced physiological perturbations impacts water use efficiency (WUE), chlorophyll florescence and photosynthesis in maize (Avramova *et al*., 2016). Current study showed reduction in shoot growth and root growth kinetics, reduction in leaf area index (LAI) and net assimilation rate (NAR) in all maize varieties (Table 1). Increased RSR in maize occurs under drought (Studer *et al*., 2017); similar observations were noted for TDI in all varieties except PG-2475 (Fig. 4). Independent studies on maize cultivars (Rung Nong 35 and Dong Dan 80) subjected to drought (35% of field capacity) compared to control irrigation (80% of field capacity) showed reduction in growth and yield in Rung Nong 35 compared to Dong Dan 80 (Anjum *et al*., 2016). Higher leaf relative water content (LRWC), proline (PL), carbohydrate and antioxidant activity (SOD, POD, CAT and GR) and lowered membrane damage accounted for better growth of Dong Dan 80 under drought compared to Rung Nong 35 (Anjum *et al*., 2016). Our observations made for TDI also showed increased LRWC, PL and glycine (Gly) accumulation, higher antioxidant activities (ASA, GSH, SOD, POD, GR and CAT) in DKC-9144 (best performing under deficit irrigation) (Supplementary Fig. S7). Reduced yield, root biomass, root length and increased root diameter has been noted in maize under salt stress (Cai *et al*., 2017). Likewise, impact of TDI on yield, root biomass and root surface area was noted for all varieties, with DKC-9144 emerged as best adapted variety (Figs. 3 and 4) in terms of WUE, cob yield and plant fitness.

Drought or water deficit negatively impacts photosynthetic performance such as net photosynthesis (P_N_), intercellular CO2 (Ci), stomatal conductance (gs) and transpiration rate (E) of crop plants (Gleason *et al*., 2019). Current study also observed deficit irrigation induced negative impact on photosynthetic performance of DKC-9144 and PG-2475 subjected to EDI alone, while EDI supplemented with PBZ improved photosynthesis related parameters (P_N_, Ci, gs and E) (Fig. 5 a-e). PBZ application was able to improve photosynthetic efficiency of both DKC-9144 and PG-2475 compared to EDI alone, probably due to delaying leaf senescence and protection of photosynthetic machinery (Pal *et al*., 2016; Mohan *et al*., 2019).

Structural equation modeling (SEM) offers a reliable tool in identification and characterization of a variable and its contribution towards a desired trait (Malaeb *et al*., 2000; Grace *et al*., 2007). Comprehensive SEM on maize root traits proposed RSA of SRs as an important trait for improving water use efficiency (WUE), shoot height (SH) and stem thickness (ST) in DKC-9144 over PG-2475 in EDI and TDI (Fig. 6 a-l). SEM was envisaged that root surface area (RSA) is an important root trait to accommodate variability, under TDI conditions SR (more specifically) and CR was negatively linked with WUE and SH (Fig. 6 a-l). Thus RSA of SRs could be a key root trait for enhancing the deficit irrigation potential of a maize variety growing in arid and semi-arid regions of the world. Additional benefit of developing a maize variety with super root trait of SRs may reduce irrigation requirement and saved water may be used to grow other crops of economic importance.

## 4.0 METHODS

### 4.1 Plant material and growth conditions

*Zea mays* L. (maize or corn) seeds of five maize varieties i.e. DDKL, DKC-9144, PG-2475, PG-2320 and Bio-9621 were procured from Agricultural Research Station Jammu, India. Seeds were tested for their germination indices. These varieties are well adapted in the climatic conditions of North India with average yield grown in irrigated and rain fed areas as follows: DDKL (5500 kg/ha), DKC-9144-9144 (6386 kg/ha), PG-2475 (3600 kg/ha), PG-2320 (3500 kg/ha) and Bio-9621 (2600 kg/ha) as per yield reports (Annual Report 2013-2014 & 2016-2017, Government of INDIA). With randomized factorial design, each variety was subjected to control irrigation regime (CN, 100% evapotranspiration demand (EVTD)), deficit irrigation (DI) with 60% EVTD with or without paclobutrazol (PBZ) application at 60 ppm in a soil drench method. Experiments over a three years period (2017, 2018 and 2019) were conducted inside a Polyhouse at the botanical garden (32.72°N 74.85°E), Botany Department, University of Jammu, India. All the environmental conditions such as photosynthetically active radiation (PAR) and temperature inside the polyhouse were monitored on regular basis. A photoperiod of 16/8 hr light/dark was maintained. Each pot (26 cm height, 25 cm diameter across top and 16 cm diameter across bottom) was filled with 7 kg of clay loam soil mixed with farmyard manure in the ratio of 3:1. Before the commencement of the deficit irrigation regimes volumetric soil water content was calculated for the soil. The following experiments were conducted:

#### *Early deficit irrigation* (EDI)

To mimic late monsoon arrival, seeds were sown at least 10 days later than their usual biological clock (1^st^ week of June, 2017, 2018 and 2019) and subjected to EDI with or without PBZ commencing from 15 days after sowing (DAS) until 35 DAS. Over a three year period, a total of 300 young maize seedlings were grown till establishment of young plants i.e. 35 DAS. In brief, for each variety 60 young maize seedlings were sown inside polyhouse until 15 DAS, with 20 plants/year in pot conditions. Each variety was subjected to four different treatments types i.e. control (CN) (100% EVTD), EDI (60% EVTD), EDI plus PBZ (EDI+PBZ) and CN plus PBZ (CN+PBZ).

#### *Terminal deficit irrigation* (TDI)

To mimic shorter rain spell, 54 DAS, plants were segregated into four groups’ viz. CN, TDI, TDI+PBZ and CN+PBZ for each maize variety. Experiments were performed on consecutive three years period (2017, 2018 and 2019) in a Polyhouse during the months of June September as a kharif crop. Each pot was fertilized with NPK twice during the life cycle of the plant at 35 DAS and 55 DAS. Over a three year period a total of 300 maize plants were grown till harvest. In brief, for each variety 60 plants were sown and fully grown until harvest over a three year period with twenty plants /year. Each variety was subjected to four different treatments CN (100% EVTD), TDI (60% EVTD), TDI plus PBZ (TDI+PBZ) and CN plus PBZ (CN+PBZ). EVTD was measured with Pen Men and Monneth equation (Mohan *et al*., 2019) using variables of the local environment on bi-weekly basis. Deficit irrigation conditions were maintained using EVTD on plant basis twice a week, such that individual plant height and leaf surface area was used to determine EVTD of a plant to induce 40% reduction in EVTD per plant.

### 4.2 Plant growth kinetics

Impact of deficit irrigation (EDI and TDI) with or without PBZ on growth kinetics were measured for shoot extension rates (SER) at 25 and 35 DAS, while root extension rates (RER) were measured for brace roots (BRs), crown roots (CRs) and seminal roots (SRs) at 25 and 35 DAS. For TDI, only shoot growth kinetics (SERt) was measured on every ten days basis commencing from TDI (at 54 DAS) until harvest (104 DAS) with or without PBZ (Yazdanbakhsh and Fisahn 2010).

Besides, growth kinetics, shoot morphometerics viz. stem height (cm), stem thickness (cm), leaf number (LN), shoot dry weight (ShDW) and root morphometerics viz. root number (RN), root length (cm), root surface areas (cm^2^) were measured for EDI and TDI. While root dry weights (RDW) were measured only for TDI.

### 4.3 Reproductive traits

Impact of TDI on successful transition of vegetative stage to reproductive stage i.e. 54 DAS to 75 DAS (stage specific for formation of reproductive organs), developmental attributes of male reproductive organs: tassel (male inflorescence) included tassel diameter (diaT, cm), length (LeT, cm), dry biomass (DBT, g), number of male flowers/plant, number of anthers/inflorescence, number of spikelets/flower, spikelet length/flower; for female reproductive organs: silk length (cm) and cob dry weight (CDW) and reproductive efforts (RE %) for tassel and ear formation were measured (Kaul *et al*., 2002).

### 4.4 Root plasticity, drought indices and physiological stress markers

Phenotypic plasticity of roots play determinant role in evaluating a plants response towards a given abiotic stressor including drought or deficit irrigation. These root plasticity indices explores the scope of plastic nature of a root system and its adaptability under changing soil environs. Root plasticity indices such that scope of plastic response (SPR, evaluates degree of roots plasticity), relative trait range (RTR, root factor (root length, root surface area) means in controlled irrigation-mean in the deficit irrigation)/absolute maximum value) and response coefficient (RC, ratio of roots mean values in control irrigation and deficit irrigation availability) and coefficient of variation (CoV, mixes variability within the variety and between environment viz. controlled irrigation and deficit irrigation). These indices were calculated for crown roots (CRs), brace roots (BRs) and seminal roots (SRs) under TDI with or without PBZ (Valladares *et al*., 2006; Zhao *et al*.,2018). Similar to plasticity indices, stress tolerance and/or susceptibility indices of a crop plant play key role in selection of genotypes resistant to various abiotic stresses including drought. To identify varietal response to deficit irrigation, drought indices namely stress susceptibility index (SSI), tolerance (TOL), mean productivity (MP), geometric mean productivity (GMP), stress tolerance index (STI), yield stability index (YSI), harmonic mean of yield (HM) and yield index (YI)) were determined (Sánchez-Reinoso *et al*., 2020). Physiological stress indices and antioxidant activities were measured as described (Pal *et al*., 2016).

### 4.5 Photosynthetic performance and genetic regulation of root traits

Photosynthetic pigments (chlorophyll *a* and *b*), leaf area index (LAI), net assimilation rate (NAR) were measured for TDI with or without PBZ (Pal *et al*., 2016; Mohan *et al*., 2019). Among five varieties, two varieties i.e. DKC-9144 and PG-2475 were selected on growth performance and water use efficiency (WUE-CDW) basis in terms of cob yield and plant height. Selected varieties were subjected to EDI with or without PBZ and photosynthetic parameters (net photosynthesis, P_N_), intercellular CO2 (Ci), stomatal conductance (gs) and leaf transpiration rate (E) were measured using LI-6400 XT infrared gas analyzer (Li-Cor, Lincoln, NE, USA) (Pal *et al*., 2016; Mohan *et al*., 2019) at two time points 25 and 35 DAS.

For quantitative real time polymerase chain reaction (qPCR), total RNA was extracted from root tissues of DKC-9144 and PG-2475 at three time points (15, 25 and 35 DAS) with or without PBZ using Nucleospin RNA plant kit (Pal *et al*., 2017) (Macherey-Nagel, Germany). cDNA synthesis and qPCR was performed as described previously (Pal *et al*., 2017) on Roche Light Cycler 96 (Roche Diagnostics, Mannheim, Germany). *Zea mays elongation factor (ZmEF α1)* was used as a house keeping gene with three independent biological replicates (*n*=3) and each biological replicate had three technical replications. Transcript levels of *ZmMel (metallothionein-like protein), ZmRTCS (gene regulating rootless crown and seminal roots), ZmRTCL (The RTCS-LIKE gene), ZmARF34 (Zea mays Auxin response factor 34)* were performed using primers published elsewhere (Xu *et al*., 2015) (Supplementary Table 3).

### 4.6 Structural equation modeling for water use efficiencies and growth performances

Structural equation modeling (SEM) tested how changes in RSA, RL and RN affected BR, CR and SR which further affected WUE, SH and ST. SEM being an advanced and robust multivariate statistical method allows for hypothesis testing of complex path-relation networks (Malaeb *et al*., 2000; Grace *et al*., 2007). AMOS, statistical software was used for the analysis of moment structures. AMOS provided overall Chi-square (χ^2^) value, together with its degrees of freedom and probability value. AMOS output gave various indices viz. Comparative fit index (CFI ≥ 0.95), Tucker-Lewis Index (TLI ≥ 0.95) and root mean square error of approximation (RMSEA ≤ 0.05) as defined by various researchers, and these indices were in acceptable range hence model fit was strengthened. SEM models considered in our study satisfied above mentioned indices. Further, we have shown the results of significant relationships in Figure 6 of root types and their traits for WUE, SH and ST under different conditions (for detailed methodology and procedure refers to SEM metadata). SEM modeling used Akaike information criterion (AIC) (Akaike 1973) and Bayesian information criterion (BIC) (Schwartz 1978), each of which was designed to penalize models with larger numbers of parameters. AIC and BIC on the number of parameters in the model:

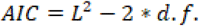

and

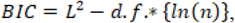

Where *n* is the sample size. Thus, models with lower values of information criteria have a better fit to a data, for a given number of parameters. When sample size is large, BIC is preferred fit statistics and for small to medium sample sizes, the AIC statistic is most commonly used (*for detailed procedure, SEM methodology*).

### 4.7 STATISTICAL ANALYSES

Experiments were arranged in a randomized factorial design (2017, 2018 and 2019). Data obtained were analyzed by ANOVA and all means were separated at the *p* < 0.05 level using the Duncan test. All calculations and data analyses were performed using the IBM SPSS 20.0 for Windows software package and the values were expressed by means and standard error (SE).

## ACKNOWLEDGEMENTS

Authors would like to thank Head, Department of Botany, University of Jammu for extending laboratory facilities and technical support and Director, Agriculture Research Station, Jammu for maize germplasm and technical assistance. The research work was financially supported by UGC-DRS-SAP-PHASE-II, New Delhi, India and CSIR-Junior Research Fellowship, Government of India to Urfan Mohammad. There are no conflicts of interest among the authors of this manuscript.

## AUTHORS’ CONTRIBUTIONS

S.P. conceived and designed the experiments; M.U., H.R.H., S.S. and M.K. performed experiments. S.P., D.V., S.B.S., S.B. and N.S.Y. performed data analysis, prepared figures and wrote the article.

## Notes

### Competing Interest Statement

The authors have declared no competing interest.

### Summary of Updates

Version has been updated with discussion and results section

## References

1. Al-Naggar, A.M.M.; Shafik, M.M.; Elsheikh, M.O.A. Putative Mechanisms of Drought Tolerance in Maize (*Zea mays* L.) via Root System Architecture Traits. Annu. Res. Rev. Biol. 2019, 1–19

2. Anjum, S.A.; Tanveer, M.; Ashraf, U.; Hussain, S.; Shahzad, B.; Khan, I.; Wang, L. Effect of progressive drought stress on growth, leaf gas exchange, and antioxidant production in two maize cultivars. Environ. Sci. Pollut. Res. 2016, 23: 17132–17141

3. Avramova, V.; Nagel, K.A.; AbdElgawad, H.; Bustos, D.; DuPlessis, M.; Fiorani, F.; Beemster, G.T. Screening for drought tolerance of maize hybrids by multi-scale analysis of root and shoot traits at the seedling stage. J. Exp. Bot. 2016, 67: 2453–2466

4. Barker, J.B.; Bhatti, S.; Heeren, D.M.; Neale, C.M.; Rudnick, D.R. Variable Rate Irrigation of Maize and Soybean in West-Central Nebraska Under Full and Deficit Irrigation. Front. Big Data. 2019, 2: 34

5. Barutcular, C.; Sabagh, A. E.; Konuskan, O.; Saneoka, H.; Yoldash, K. M. Evaluation of maize hybrids to terminal drought stress tolerance by defining drought indices. J. Exper. Biol. Agricul. Sci. 2016, 4(6), 610–616.

6. Begcy, K.; Nosenko, T.; Zhou, L.Z.; Fragner, L.; Weckwerth, W.; Dresselhaus, T. Male sterility in maize after transient heat stress during the tetrad stage of pollen development. Plant physiol. 2019, 683–700

7. Borrás, L.; Vitantonio-Mazzini, L.N. Maize reproductive development and kernel set under limited plant growth environments. J. Exp. Bot. 2018, 3235–3243

8. Cai, R.; Dai, W.; Zhang, C.; Wang, Y.; Wu, M.; Zhao, Y.;… Cheng, B. The maize WRKY transcription factor ZmWRKY17 negatively regulates salt stress tolerance in transgenic Arabidopsis plants. Planta. 2017, 246: 1215–1231

9. Challinor, A.J.; Koehler, A.K.; Ramirez-Villegas, J.; Whitfield, S.; Das, B. Current warming will reduce yields unless maize breeding and seed systems adapt immediately. Nat. Clim. Change. 2016, 6: 954–958

10. Comas, L.; Becker, S.; Cruz, V.M.V.; Byrne, P.F.; Dierig, D.A. Root traits contributing to plant productivity under drought. Front. Plant Sci. 2013, 4: 442

11. David, L. Maize vulnerable to drought. Nature. 2014, 137 https://doi.org/10.1038/509137d

12. Davies, W.J.; Bennett, M.J. Achieving more crop per drop. Nat. Plants. 2015, 1: 1–2

13. Deng, K.; Yang, S.; Ting, M.; Tan, Y.; He, S. Global monsoon precipitation: Trends, leading modes, and associated drought and heat wave in the Northern Hemisphere. J. Clim. 2018, 6947–6966

14. Dowswell, C. Maize in the third world. CRC Press. 2019.

15. Gleason, S. M.; Cooper, M.; Wiggans, D. R.; Bliss, C. A.; Romay, M. C.; Gore, M. A.;… Comas, L. H. Stomatal; conductance, xylem water transport, and root traits underpin improved performance under drought and well-watered conditions across a diverse panel of maize inbred lines. Field Crops Res. 2019, 234, 119–128.

16. Grace, J.B.; Michael, A. T.; Smith, M.D.; Seabloom, E.; Andelman, S.J.; Meche, G.’;… Knops, J. Does species diversity limit productivity in natural grassland communities?. Ecol. Lett. 2007, 10: 680–689

17. Guo, J.; Li, C.; Zhang, X.; Li, Y.; Zhang, D.; Shi, Y.;… Wang, T. Transcriptome and GWAS analyses reveal candidate gene for seminal root length of maize seedlings under drought stress. Plant Sci. 2020, 292: 110380

18. He, Q.; Zhou, G.; Lü, X.; Zhou, M. Climatic suitability and spatial distribution for summer maize cultivation in China at 1.5 and 2.0° C global warming. Sci. Bull. 2019, 64: 690–697

19. http://ficci.in/ficci-in-news-page.asp?nid=14261

20. https://mausam.imd.gov.in/

21. https://www.farmer.gov.in/M_cropstaticsmaize.aspx

22. Hund, A.; Ruta, N.; Liedgens, M. Rooting depth and water use efficiency of tropical maize inbred lines, differing in drought tolerance. Plant Soil. 2009, 318: 311–325

23. Jägermeyr, J.; Frieler, K. Spatial variations in crop growing seasons pivotal to reproduce global fluctuations in maize and wheat yields. Sci. Adv. 2018, 4: eaat4517

24. Kaul, V.; Sharma, N.; Koul, A.K. Reproductive effort and sex allocation strategy in *Commelina benghalensis* L., a common monsoon weed. Bot. J. Linn. Soc. 2002, 140: 403–413

25. Klein, S.P.; Schneider, H.M.; Perkins, A.C.; Brown, K.M.; Lynch, J.P. Multiple integrated root phenotypes are associated with improved drought tolerance. Plant Physiol. 2020, 1011–1025.

26. Laitonjam, N.; Singh, R.; Feroze, S.M. Effect of Pre-Monsoon rainfall on maize yield in Manipur. Econ. Affairs. 2018, 413–417

27. Lesk, C.; Rowhani, P.; Ramankutty, N. Influence of extreme weather disasters on global crop production. Nature. 2016, 529: 84–87

28. Malaeb, Z.A.; Summers, J.K.; Puge, sek. B.H. Using structural equation modelling to investigate relationships among ecological variables. Environ. Ecol. Stat. 2000, 7: 93–111

29. Manivasagam, V.S.; Nagarajan, R. Rainfall and crop modeling-based water stress assessment for rainfed maize cultivation in peninsular India. Theor. Appl. Climatol. 2018, 132, 529–542

30. Menkir, A.; Crossa, J.; Meseka, S.; Bossey, B.; Muhyideen, O.; Riberio, P.F.;… Haruna, A. Staking Tolerance to Drought and Resistance to a Parasitic Weed in Tropical Hybrid Maize for Enhancing Resilience to Stress Combinations. Front. Plant Sci. 2020, 11: 166

31. Mohan, R.; Kaur, T.; Bhat, H.A.; Khajuria, M.; Pal, S.; Vyas, D. Paclobutrazol Induces Photochemical Efficiency in Mulberry (*Morus alba* L.) Under Water Stress and Affects Leaf Yield Without Influencing Biotic Interactions. J. Plant Growth Regul. 2019, 39: 205–215

32. Murari, K.K.; Jayaraman, T.; Swaminathan, M. Climate change and agricultural suicides in India. Proc. Natl. Acad. Sci. USA. 2018, 115: E115–E115

33. Oury, V.; Tardieu, F.; Turc, O. Ovary apical abortion under water deficit is caused by changes in sequential development of ovaries and in silk growth rate in maize. Plant Physiol. 2016, 171: 986–996

34. Pal, S.; Kisko, M.; Dubos, C.; Lacombe, B.; Berthomieu, P.; Krouk, G.; Rouached, H. Transdetect identifies a new regulatory module controlling phosphate accumulation. Plant Physiol. 2017, 175: 916–926

35. Pal, S.; Zhao, J.; Khan, A.; Yadav, N.S.; Batushansky, A.; Barak, S.;… Rachmilevitch, S. Paclobutrazol induces tolerance in tomato to deficit irrigation through diversified effects on plant morphology, physiology and metabolism. Sci. Rep. 2016, 6: 39321

36. Parmar, H.C.; Zala, Y.C.; Mor, V.B. Economic Analysis of Rainfed Maize Production in Central Gujarat, India. Int J Curr Microbiol. Appl. Sci. 2019, 2420–2428

37. Pires, M. V.; de Castro, E.M.; de Freitas, B. S. M.; Lira, J. M. S.; Magalhães, P. C.; Pereira, M. P. Yield-related phenotypic traits of drought resistant maize genotypes. Env. Exp. Bot. 2020, 171, 103962.

38. Sah, R.P.; Chakraborty, M.; Prasad, K.; Pandit, M.; Tudu, V.K.; Chakravarty, M.K.;… Moharana, D. Impact of water deficit stress in maize: Phenology and yield components. Sci. Rep. 2020, 10: 1–15

39. Sánchez-Reinoso, A.D.; Ligarreto-Moreno, G.A.; Restrepo-Díaz, H. Evaluation of drought indices to identify tolerant genotypes in common bean bush (*Phaseolus vulgaris* L.). J. Integr. Agric. 2020, 19: 99–107

40. Sebastian, J.; Yee, M.C.; Viana, W.G.; Rellán-Álvarez, R.; Feldman, M.; Priest, H.D.;… Mockler, T.C. Grasses suppress shoot-borne roots to conserve water during drought. Proc. Natl. Acad. Sci. USA. 2016, 8861–8866

41. Seneviratne, S.I. Climate science: Historical drought trends revisited. Nature. 2012, 491: 338–339

42. Sharma, S.; Mujumdar, P. Increasing frequency and spatial extent of concurrent meteorological droughts and heat waves in India. Sci. Rep. 2017, 7: 1–9

43. Sloat, L.L.; Davis, S.J.; Gerber, J.S.; Moore, F.C.; Ray, D.K.; West, P.C.; Mueller, N.D. Climate adaptation by crop migration. Nat. Commun. 2020, 11: 1–9

44. Studer. C.; Hu, Y.; Schmidhalter, U. Interactive effects of N-, P-and K-nutrition and drought stress on the development of maize seedlings. Agriculture. 2017, 7: 90

45. Thatcher, S.R.; Danilevskaya, O.N.; Meng, X.; Beatty, M.; Zastrow-Hayes, G.; Harris, C.;… Li, B. Genome-wide analysis of alternative splicing during development and drought stress in maize. Plant Physiol. 2016, 170: 586–599

46. Thomas, J.; Prasannakumar, V. Temporal analysis of rainfall (1871–2012) and drought characteristics over a tropical monsoon-dominated State (Kerala) of India. J. Hydrol. 2016, 266–280.

47. Trout. T.J.; Manning, D.T. An Economic and Biophysical Model of Deficit Irrigation. Agron. J. 2019, 111: 3182–3193

48. Turc, O.; Tardieu, F. Drought affects abortion of reproductive organs by exacerbating developmentally driven processes via expansive growth and hydraulics. J. Exp. Bot. 2018, 3245–3254

49. Valladares, F.; SANCHEZ-GOMEZ, D.; Zavala, M.A. Quantitative estimation of phenotypic plasticity: bridging the gap between the evolutionary concept and its ecological applications. J. Ecol. 2006, 1103–1116.

50. Wang, N.; Li, L.; Gao, Ww. WY.; Yong, Hj. WJ. Transcriptomes of early developing tassels under drought stress reveal differential expression of genes related to drought tolerance in maize. J. Integr. Agric. 2018, 17: 1276–88.

51. Westgate, M.E. Seed formation in maize during drought. Physiology and determination of crop yield, 1994, pp. 361–364

52. Xu, C.; Tai, H.; Saleem, M.; Ludwig, Y.; Majer, C.; Berendzen, K.W.;… Hochholdinger, F. Cooperative action of the paralogous maize lateral organ boundaries (LOB) domain proteins RTCS and RTCL in shoot-borne root formation. New Phytol. 2015, 207: 1123–1133

53. Xu, C.; Zhao, H.; Zhang, P.; Wang, Y.; Huang, S.; Meng, Q.; Wang, P. Delaying wheat seeding time and maize harvest improved water use efficiency in a warm temperature continental monsoon climate. Agronomy J. 2018. 1420–1429.

54. Yazdanbakhsh, N.; Fisahn, J. Analysis of *Arabidopsis thaliana* root growth kinetics with high temporal and spatial resolution. Ann. Bot. 2010, 783–791

55. Zaveri, E.; Russ, J.; Damania, R. Rainfall anomalies are a significant driver of cropland expansion. Proc. Natl. Acad. Sci. USA. 2020, 117: 10225–10233.

56. Zhan, A.; Schneider, Lynch, J.P. Reduced lateral root branching density improves drought tolerance in maize. Plant physiol. 2015, 1603–1615

57. Zhang, H.; Li, Y.; Zhu, J.K. Developing naturally stress-resistant crops for a sustainable agriculture. Nat. Plants. 2018, 4: 989–996

58. Zhao, J.; Sykacek, P.; Bodner, G.; Rewald, B. Root traits of European *Vicia faba* cultivars—Using machine learning to explore adaptations to agroclimatic conditions. Plant Cell Environ. 2018, 1984–1996

59. Zhuang, Y.; Ren, G.; Yue, G.; Li, Z.; Qu, X.; Hou, G.;… Zhang, J. Effects of water-deficit stress on the transcriptomes of developing immature ear and tassel in maize. Plant Cell Rep. 2007, 26: 2137–2147

